# Multiplex genetic engineering exploiting pyrimidine salvage pathway-based self-encoded selectable markers

**DOI:** 10.1101/2020.01.16.908764

**Authors:** Lukas Birštonas, Alex Dallemulle, Manuel S. López-Berges, Ilse D. Jacobsen, Martin Offterdinger, Beate Abt, Maria Straßburger, Ingo Bauer, Oliver Schmidt, Bettina Sarg, Herbert Lindner, Hubertus Haas, Fabio Gsaller

**Author notes:** Department of Genetics, University of Córdoba, Córdoba, Spain (Manuel S. López-Berges).

## Abstract

Selectable markers are indispensable for genetic engineering, yet their number and variety is limited. Most selection procedures for prototrophic cells rely on the introduction of antibiotic resistance genes. New minimally invasive tools are needed to facilitate sophisticated genetic manipulations. Here, we characterized three endogenous genes in the human fungal pathogen *Aspergillus fumigatus* for their potential as markers for targeted genomic insertions of DNAs-of-interest (DOIs). Since these genes are involved in uptake and metabolization of pyrimidines, resistance to the toxic effects of prodrugs 5-fluorocytosine and 5-fluorouracil can be used to select successfully integrated DOIs. We show that DOI integration, resulting in the inactivation of these genes, caused no adverse effects with respect to nutrient requirements, growth or virulence. Beside the individual use of markers for site-directed integration of reporter cassettes including the 17-kb penicillin biosynthetic cluster, we demonstrate their sequential use inserting three fluorescent protein encoding genes into a single strain for simultaneous multicolor localization microscopy. In addition to *A. fumigatus*, we validated the applicability of this novel toolbox in *Penicillium chrysogenum* and *Fusarium oxysporum*.

Enabling multiple targeted insertions of DOIs without the necessity for exogenous markers, this technology has the potential to significantly advance genetic engineering.

## INTRODUCTION

Genetic engineering commonly involves the introduction of DNA-of-interest (DOI) into a target cell followed by its integration into the genome. However, the efficiency of such genetic transformations is typically very limited. Therefore, selection of successfully modified cells usually involves co-transformation of selectable marker genes together with the DOI to allow growth under selective conditions. Widely used marker cassettes either compensate for the inability to synthesize vital metabolites (auxotrophic selection markers) or confer resistance to growth inhibitory compounds such as antibiotics (dominant selectable markers) (Su et al. 2012). While auxotrophic markers are restricted to auxotrophic recipients, dominant selectable markers can be used for virtually any prototrophic recipient cell that is susceptible to the antibiotic used for selection. Hence, dominant selectable markers play a crucial role in the genetic manipulation of most wild-type cells. Examples of selectable markers commonly employed for the genetic engineering of fungi include hygromycin B, phleomycin, pyrithiamine, kanamycin and nourseothricin resistance conferring genes (Punt and van den Hondel 1992; Kubodera et al. 2002; Kück and Hoff 2006). Their expression allows growth in the presence of the corresponding antibiotic, which classifies them as positive selectable markers. Negative selectable markers, in contrast, inhibit growth of the target cells during selective conditions. In addition to the herpes simplex virus-1 thymidine kinase gene (Wigler et al. 1977; Borrelli et al. 1988), genes encoding cytosine deaminase (CD) and uracil phosphoribosyltransferase (UPRT) have been employed as negative selectable markers in diverse organisms (Mullen et al. 1992; Fox et al. 1999; van der Geize et al. 2008; Orr et al. 2012; Shi et al. 2013). The presence of functional gene copies providing CD and UPRT activity renders host cells susceptible to prodrugs 5-fluorocytosine (5FC) and 5-fluorouracil (5FU). CD and UPRT, both enzymes of the pyrimidine salvage pathway, convert 5FC and 5FU into 5-fluorouridine monophosphate (5FUMP) (Fig. 1a). Further metabolization of 5FUMP into toxic ribo- and deoxyribonucleotides blocks cellular growth (Vermes et al. 2000).

**Fig 1.**
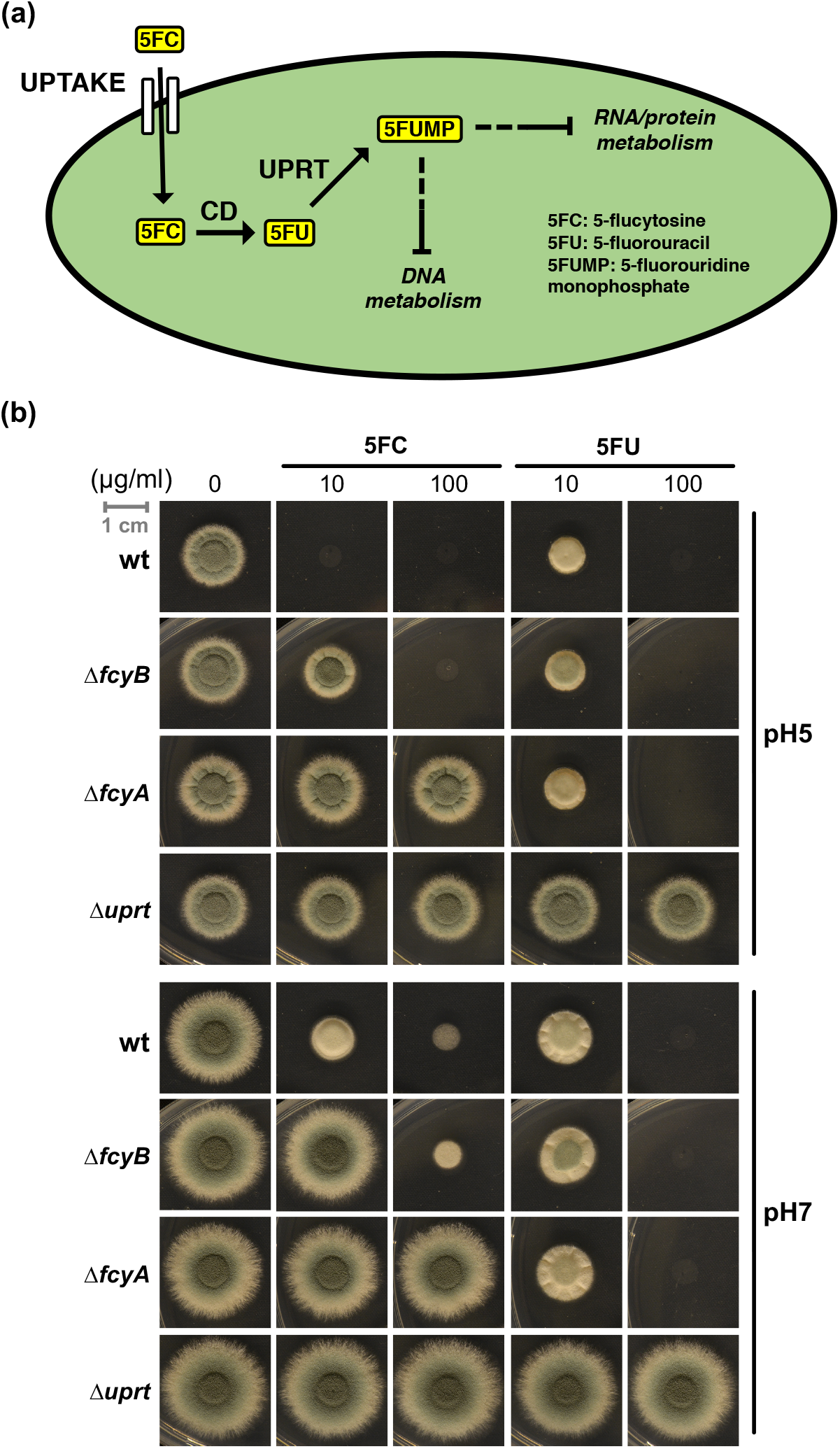
Metabolization of 5FC and associated genetic factors in *A. fumigatus*. (a) After uptake, 5FC is converted to 5FU by the enzyme cytosine deaminase (CD). Subsequently, 5FU is phosphoribosylated to 5FUMP by uracil phosphoribosyltransferase (UPRT). 5FUMP is further metabolized into RNA or DNA nucleotides that interfere with DNA, RNA as well as protein metabolism (Vermes et al. 2000). (b) Inactivation of genes encoding uptake (Δ*fcyB*), CD (Δ*fcyA*) and UPRT (Δ*uprt*) activity in *A. fumigatus* leads to different degrees of 5FC and 5FU resistance. For plate growth-based susceptibility testing, strains were point inoculated on solid AMM containing different levels of 5FC and 5FU. Images were acquired after 48 h incubation at 37 °C. Dashed lines indicate several enzymatic steps.

Here, we characterized the pyrimidine salvage pathway in the human fungal pathogen *Aspergillus fumigatus* and present its application for fungal genetic engineering. In the described technology, endogenous genes encoding 5FC uptake, CD and UPRT serve as counterselectable markers for targeted, genomic introduction of multiple DOIs. Homologous recombination-driven replacement of marker genes by DOIs results in their inactivation, which can be selected *via* 5FC/5FU resistance. In addition to the individual use (e.g. integration of reporter cassettes as well as the 17-kb penicillin biosynthetic cluster), the potential sequential use of the three loci is demonstrated by the insertion of three different fluorescent protein-encoding genes for multicolor imaging of three cellular compartments. Demonstrating its versatile applicability, the described technology was implemented in the industrial work-horse *Penicillium chrysogenum* and the plant pathogen *Fusarium oxysporum*.

## RESULTS

### Cytosine deaminase FcyA and uracil phosphoribosyltransferase Uprt are crucial for the metabolic activation of 5FC in *Aspergillus fumigatus*

Metabolization of 5FC has been well-studied in the model yeast *Saccharomyces cerevisiae:* 5FC is converted by the CD Fcy1p to 5FU (Polak et al. 1976; Whelan 1987) and, subsequently, phosphoribosylated to 5FUMP by the UPRT Fur1p (Kern et al. 1990). Inactivation of each of these steps resulted in 5FC resistance, whereby inactivation of Fur1p also conferred 5FU resistance (Kern et al. 1990). Orthologous proteins from *S. cerevisiae* (Fcy2p), *A. fumigatus* and *Aspergillus nidulans* (FcyB), respectively, have been identified as major cellular 5FC importers (Paluszynski et al. 2006; Vlanti and Diallinas 2008; Gsaller et al. 2018).

Among other fungal species, *A. fumigatus* is susceptible to 5FC (Te Dorsthorst et al. 2004; te Dorsthorst et al. 2005) and is therefore anticipated to harbor genes encoding CD and UPRT activities in addition to 5FC uptake. BLASTP-based *in silico* predictions revealed *A. fumigatus* FcyA (AFUB_005410) and Uprt (AFUB_053020) as putative orthologs of yeast Fcy1p and Fur1p, respectively. To analyze their role in 5FC as well as 5FU metabolism, we inactivated *fcyA* and *uprt* in the *A. fumigatus* strain A1160P+ (Fraczek et al. 2013), termed wt here, using hygromycin and phleomycin resistance-based deletion cassettes. Due to the interdependency of 5FC activity and environmental pH (Te Dorsthorst et al. 2004; Gsaller et al. 2018), we investigated the contribution of both enzymes, as well as FcyB, to 5FC and 5FU activity at both pH5 and pH7.

Plate growth-based susceptibility testing revealed that, similar to previous work (Gsaller et al. 2018), 5FC levels ≥ 10 μg/ml blocked wt growth at pH5, while 100 μg/ml 5FC were required at pH7 (Fig. 1b). Although FcyB is the major 5FC uptake protein, at 100 μg/ml 5FC Δ*fcyB* was not able to grow at pH5 and showed severe growth inhibition at pH7. In contrast to Δ*fcyB*, Δ*fcyA* and Δ*uprt* displayed full resistance to 5FC up to 100 μg/ml, regardless of the pH. Furthermore, 100 μg/ml 5FU blocked growth of wt, Δ*fcyA* and Δ*fcyB* at pH5 as well as pH7, while Δ*uprt* was resistant.

Our data confirm the role of FcyB as major 5FC cellular importer but indicate the presence of additional uptake mechanisms. Similar to the orthologous proteins in *S. cerevisiae*, our findings reveal the essential roles of FcyA and Uprt for 5FC activity and of Uprt for metabolic activation of 5FU in *A. fumigatus*.

### Self-encoded loci *fcyB, fcyA* and *uprt* can be used for 5FC/5FU-based transformation selection

Lack of FcyB, FcyA (CD activity) or Uprt (UPRT activity) confers resistance to 5FC (Δ*fcyB*, Δ*fcyA* and Δ*uprt*) or 5FU (Δ*uprt*) (Fig. 1b), which suggested the utilization of the encoding genes as counterselectable markers for positive selection of cells with targeted integration of DOIs. Moreover, the different degrees in 5FC resistance observed for Δ*fcyB* and Δ*fcyA* indicated that 5FC can be used for selection of loss of FcyB at low 5FC concentrations (10 μg/ml) and loss of FcyA at high 5FC levels (100 μg/ml) (Fig. 1b). Selection for loss of Uprt was carried out at 100 μg/ml 5FU.

For proof-of-principle, both green fluorescent protein (GFP) and ß-galactosidase (LacZ) expression cassettes were used to replace *fcyB*, *fcyA* as well as *uprt*. To achieve homologous recombination-mediated replacement of these loci with the reporter cassettes, approximately 1 kb 5’- and 3’-non-translated regions (NTRs) of the respective gene were linked to each cassette *via* fusion PCR (Fig. 2a). The yielding knock-in constructs were transformed into wt protoplasts, which underwent selection for resistance to 5FC and 5FU (see above; Fig. 2b). Southern blot analyses confirmed site-specific integration of the DOIs into each of the three loci (Fig. S1). In agreement, all knock-in strains displayed resistance phenotypes according to the respective mutation in the pyrimidine salvage pathway (compare Fig. 1b and Supplementary Fig. 2).

**Fig 2.**
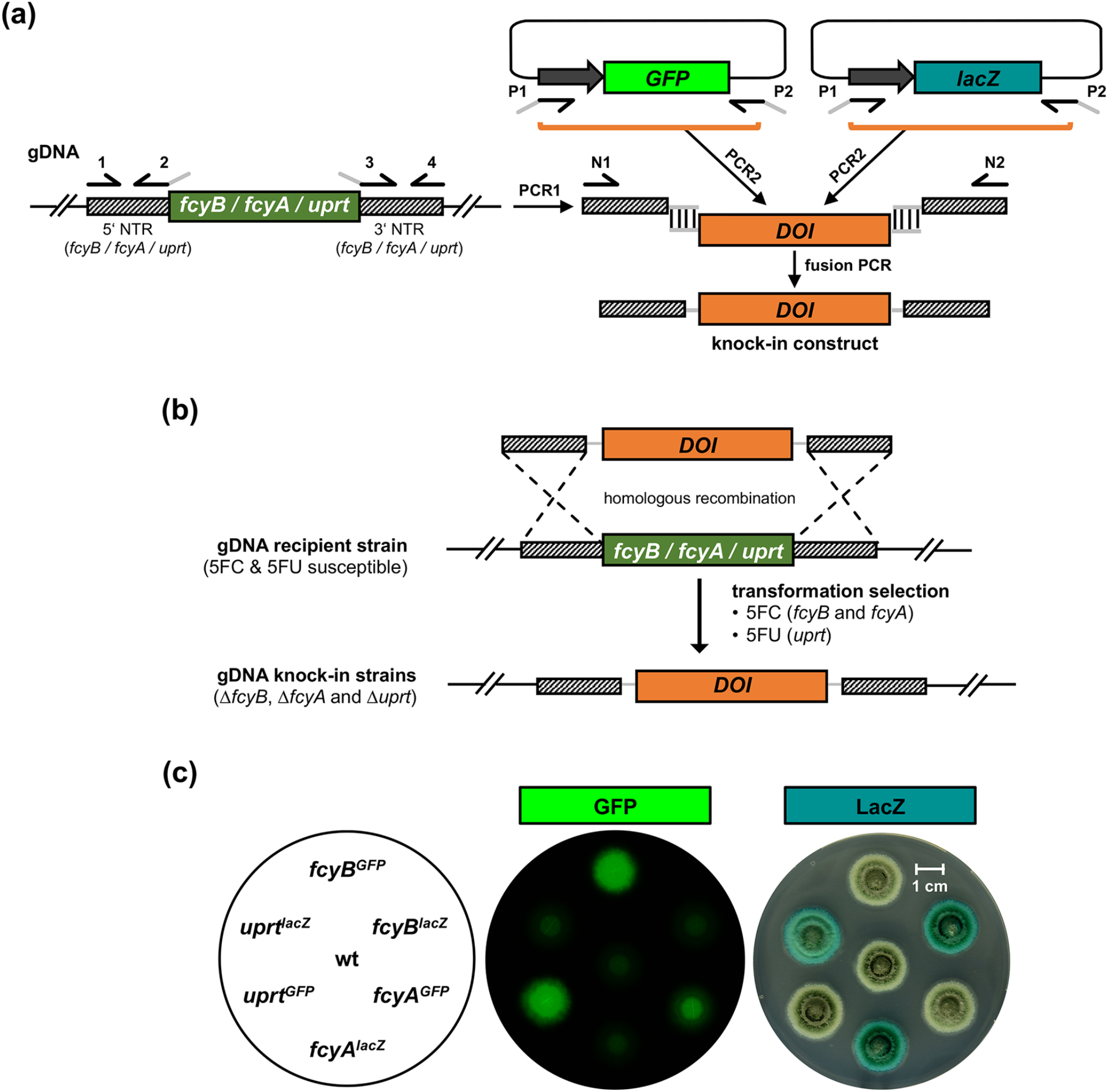
Replacement of self-encoded selectable markers *fcyB, fcyA* and *uprt* by DOIs. (a) Scheme of the generation of knock-in constructs. 5’ and 3’ NTR (PCR1) of the respective loci as well as the DOIs (PCR2; *GFP* or *lacZ* reporter cassette) are amplified from genomic DNA (gDNA) and plasmid DNA, respectively. Both NTRs and DOIs contain overlapping DNA (gray line) for subsequent connection via fusion PCR, yielding the knock-in constructs. (b) Double crossover homologous recombination-based replacement of *fcyB, fcyA* or *uprt* by DOIs. Transformation selection was carried out using 5FC (*fcyB* and *fcyA* locus) or 5FU (*uprt* locus). (c) Visualization of *GFP* as well as *LacZ* expression in the corresponding knock-in strains after incubation on solid AMM for 48h at 37°C.

Fluorescence imaging and ß-galactosidase staining confirmed functionality of the knock-in cassettes (Fig. 2c). To analyze the transformation efficiency using individual selectable marker genes, we employed the corresponding LacZ knock-in constructs for each locus. For *fcyB, fcyA* and *uprt*, 10, 27 and 13 transformants, respectively, were recovered on the corresponding selective media (Supplementary Fig. 3). Out of these 10 (100%), 26 (97%) and 12 (92%) displayed LacZ activity. Southern blot analysis of 10 LacZ-positive transformants for each locus confirmed correct integrations.

Taken together, 5FC- and 5FU-mediated selection allowed replacement of each of the three loci by either *GFP*- or *lacZ*-expression cassettes, which demonstrates the suitability of *fcyB, fcyA* and *uprt* as selectable markers for targeted, integrative transformation in *A. fumigatus*.

### *fcyB, fcyA* and *uprt* can be consecutively used for multiple genomic integrations

Due to the fact that inactivation of *fcyB* and *fcyA* led to different levels of resistance to 5FC and inactivation of *uprt* caused resistance to 5FU (Fig. 1b), we investigated if these marker genes can be sequentially employed for transformation selection in a single strain. As an exemplary application, we aimed to generate a strain expressing three fluorescent proteins for multicolor imaging: GFP (sGFP), red fluorescent protein (RFP, mKate2) and blue fluorescent protein (BFP, mTagBFP2).

The pursued strategy and order of markers used for selection was based on the following aspects (Fig. 1b): (i) in contrast to wt, Δ*fcyB* can grow in the presence of 10 μg/ml 5FC at pH5; (ii) in contrast to Δ*fcyB, ΔfcyA* can grow at 100 μg/ml 5FC, which allows discrimination of Δ*fcyBΔfcyA* from Δ*fcyB;* (iii) Δ*fcyB* and Δ*fcyA* are still able to import and metabolize 5FU, which is expected to allow discrimination of Δ*fcyB*Δ*fcyA* and Δ*fcyB*Δ*fcyA*Δ*uprt* in the presence of 100 μg/ml 5FU.

In the first step, we integrated an expression cassette encoding mKate2 carrying a C-terminal peroxisomal targeting sequence (PTS1, tripeptide SKL) (Olivier and Krisans 2000) in the *fcyB* locus, yielding strain *RFP^PER^* (Δ*fcyB::mKate2^PER^*). In this strain, a second expression cassette encoding sGFP containing an N-terminal mitochondrial targeting sequence derived from citrate synthase (Min et al. 2010) was targeted into the *fcyA* locus, yielding strain *RFP^PER^GFP^MIT^* (Δ*fcyB::mKate2^PER^*Δ*fcyA::sGFP^MIT^*). In the last step, an expression cassette encoding mTagBFP2 with expected cytoplasmic localization was integrated into the *uprt* yielding strain *RFP^PER^GFP^MIT^BFP^CYT^* (Δ*fcyB::mKate2^PER^*Δ*fcyA::sGFP^MIT^*Δ*uprt::mTagBFP2^CYT^*). Multicolor laser scanning confocal microscopy confirmed expression of all three fluorescent proteins in *RFP^PER^GFP^MIT^BFP^CYT^* and localization to distinct subcellular compartments (Fig. 3 and Supplementary Fig. 4). A noteworthy finding was that the lack of FcyB, FcyA and Uprt (strain *RFP^PER^GFP^MIT^BFP^CYT^*) affected neither growth nor virulence of *A. fumigatus* in a murine model of invasive aspergillosis (Supplementary Fig. 5).

**Fig 3.**
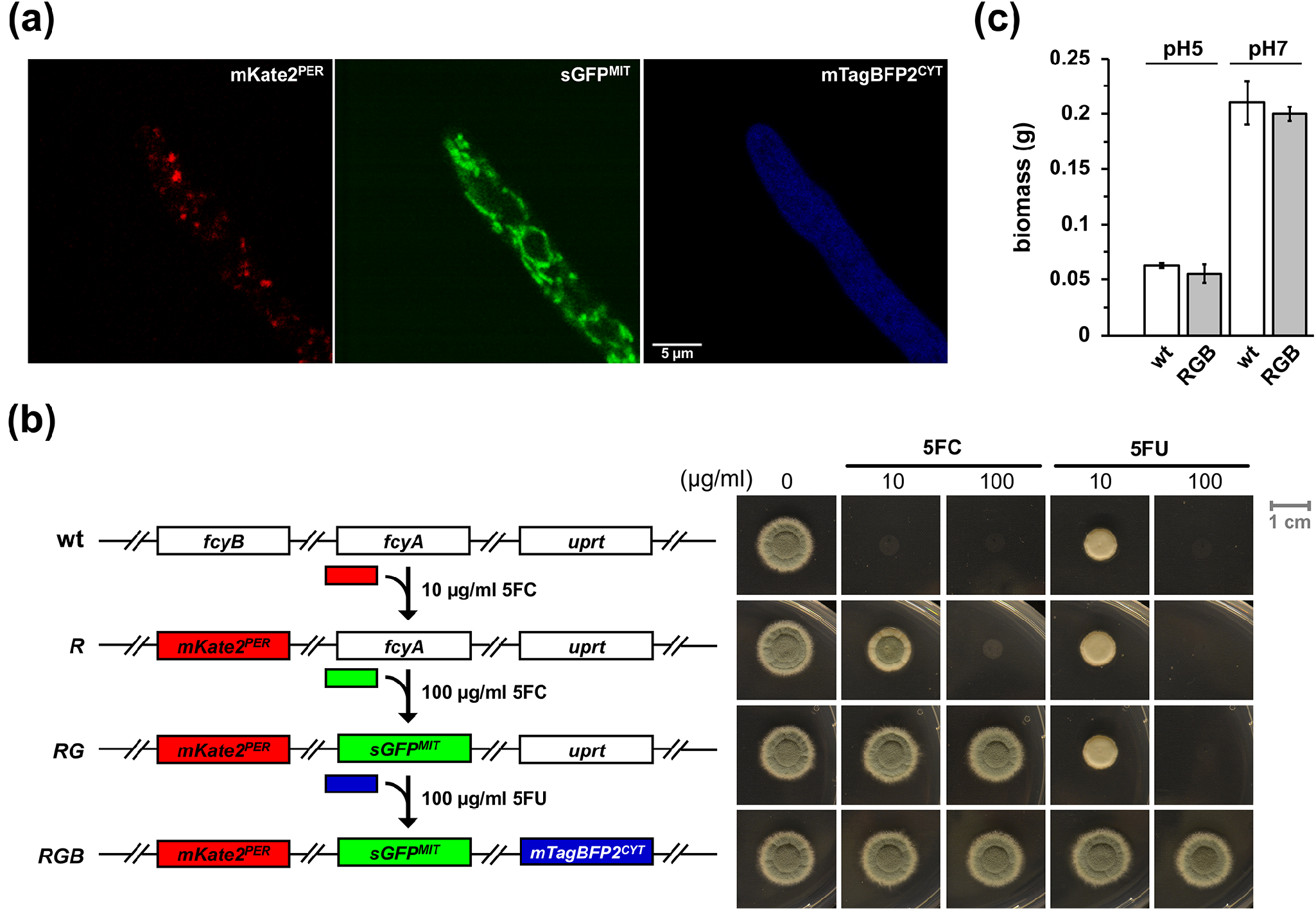
Multicolor imaging and phenotypic analysis of *RFP^PER^GFP^MIT^BFP^CYT^ (RGB)* expressing three fluorescent proteins with distinct subcellular localization. (a) Using laser scanning confocal microscopy, the expression of peroxisomal RFP, mitochondrial GFP and cytoplasmic BFP was monitored in *RGB* after incubation for 20 h in liquid AMM at 30 °C. (b) Scheme illustrating the sequential integration of fluorescent proteins (left) and 5FC/5FU resistance profiles (right) of *RFP^PER^* (*R*), *RFP^PER^GFP^MIT^ (RG)* and *RGB* after incubation on solid AMM for 48 h. (c) Biomass production (dry weight) of *RGB* and wt. Liquid cultures were incubated for 20 h at pH5 and pH7. The data illustrate the mean of biological triplicates. Error bars indicate the standard deviation. p-values were calculated by Student’s T-test (two-tailed, unpaired): 0.38 and 0.32 for pH5 and pH7, respectively (reference: wt).

Collectively, these data demonstrate the feasibility of sequential use of self-encoded selection markers *fcyB, fcyA* and *uprt* for integration of up to three DOIs without adverse effects on *A. fumigatus* growth and virulence.

### Self-encoded markers can be used for the integration of biotechnologically-relevant, large DNA fragments

Fungi play important roles as cell factories in food industry as well as medicine. Pursuing a potential biotechnological approach using the described selection method, we tested if we can integrate the 17-kb penicillin biosynthetic cluster (PcCluster) of *P. chrysogenum* into the genome of *A. fumigatus*. Therefore, a knock-in plasmid was constructed comprising the PcCluster as well as 5’- and *3’-fcyB* NTR. Linearization of the plasmid with *PmeI* allowed double crossover homologous recombination-mediated replacement of *fcyB* by the PcCluster, as illustrated in Fig. 4a and Supplementary Fig. 6.

**Fig 4.**
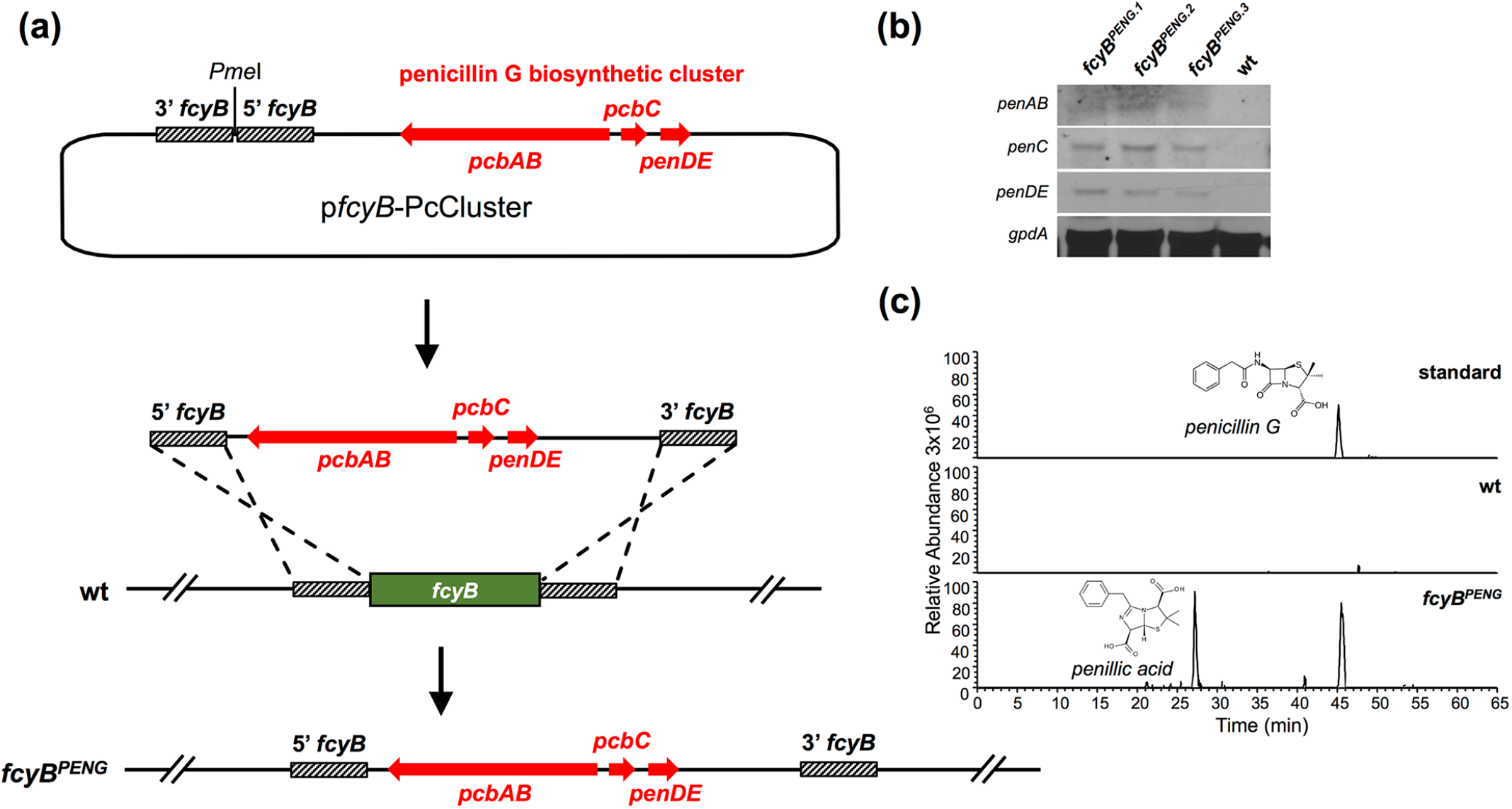
Genomic insertion of the PcCluster transformed *A. fumigatus* into a penicillin G producer. (a) To facilitate genomic integration of the PcCluster at the *fcyB* locus, the plasmid p*fcyB*-PcCluster comprising the respective DOI (17 kb) as well as *fcyB* 5’ and 3’ NTRs was generated. Linearization of this plasmid with *PmeI* allows homologous recombination-based replacement of *fcyB* coding sequence with DNA containing the PcCluster. (b) Expression of functional *pcbAB, pcbC* and *penDE* was monitored in three independent transformants using Northern blot analysis *(gpdA* was used as reference). (c) LC-MS/MS extracted ion chromatograms of penicillin G (peak at 45 min) and its degradation product penillic acid (peak at 27 min) in the culture supernatant of *fcyB^PENG^* after shaking incubation for 48 h at 25 °C. wt served as negative control.

Subsequent to transformation of this construct in wt (selection: 10 μg/ml 5FC, pH5), we validated its site-specific integration at the *fcyB* locus (strain *fcyB^PENG^*, Supplementary Fig. 1). Northern blot analysis confirmed expression of penicillin biosynthetic genes *pcbAB, pcbC* and *penDE* in three transformants (Fig. 4b). Concomitant to successful expression of this heterologous gene cluster in *A. fumigatus*, we detected penicillin G and its degradation product penillic acid in culture supernatants using nanoLC mass spectrometry (Fig. 4c). Both substances have the same molecular weight of 335.1060 g/mol but can be differentiated due to structures pecific fragmentation during MS/MS (Supplementary Fig. 6b).

Taken together, these results demonstrate that pyrimidine salvage-based selectable markers can be applied for the integration of large-size DNA fragments including whole gene clusters.

### Pyrimidine salvage based-selectable markers can be utilized in *Penicillium chrysogenum* and *Fusarium oxysporum*

To identify encoded CD and UPRT activities in further fungal species, we searched for *A. fumigatus* FcyA and Uprt orthologs in biotechnologically *(Aspergillus niger, Aspergillus oryzae, P. chrysogenum, Komagataella phaffii* (previously *Pichia pastoris) S. cerevisiae* and *Trichoderma reesei)* and pathologically relevant fungal species *(Candida albicans, Cryptococcus neoformans* and *F. oxysporum). In silico* inspection of the annotated genomes of these species revealed that 8 out of the 10 species analyzed harbor a putative ortholog of *A. fumigatus* FcyA and that all species possess a putative ortholog of *A. fumigatus* Uprt with overall sequence identities ≥ 40% (Supplementary Table 1). Notably, all species encoding an FcyA ortholog were also found to encode an FcyB ortholog. The genetic coupling of these two features might indicate that their main function is the utilization of extracellular pyrimidines.

To confirm CD and UPRT activities in the species analyzed, we monitored 5FC and 5FU susceptibility profiles following a broth microdilution-based method according to EUCAST (Subcommittee on Antifungal Susceptibility Testing of the 2008). In agreement with our homology search, species with predicted FcyA orthologs (CD activity) were susceptible to 5FC, while those without were resistant to the drug (Supplementary Table 2). All strains were susceptible to 5FU, which is in accordance with *in silico* predicted Uprt orthologs (UPRT activity).

In the next step we tested the applicability of the described selection strategy in *P. chrysogenum* and *F. oxysporum*. In agreement with the genomic data and 5FC/5FU susceptibility, *P. chrysogenum* expresses both CD (Pc-FcyA, EN45_039280) and UPRT (Pc-Uprt, EN45_060980), while *F. oxysporum* lacks CD but expresses UPRT (Fo-Uprt, FOXG_01418). Employing the same protocol as used for *A. fumigatus* enabled the integration of GFP expression cassettes flanked by 5’ and 3’ NTR of the respective *P. chrysogenum* genes in both the *Pc-fcyA* and the *Pc-uprt* loci. In *F. oxysporum*, the same strategy enabled the targeting of a GFP expression cassette at the *Fo-uprt* locus. The presence and functionality of the GFP reporters was visualized as described above (Fig. 5). As observed for *A. fumigatus* knock-in mutants, resistance profiles of *P. chrysogenum* and *F. oxysporum* knock-in strains were in accordance with the absence of individual salvage activities.

**Fig 5.**
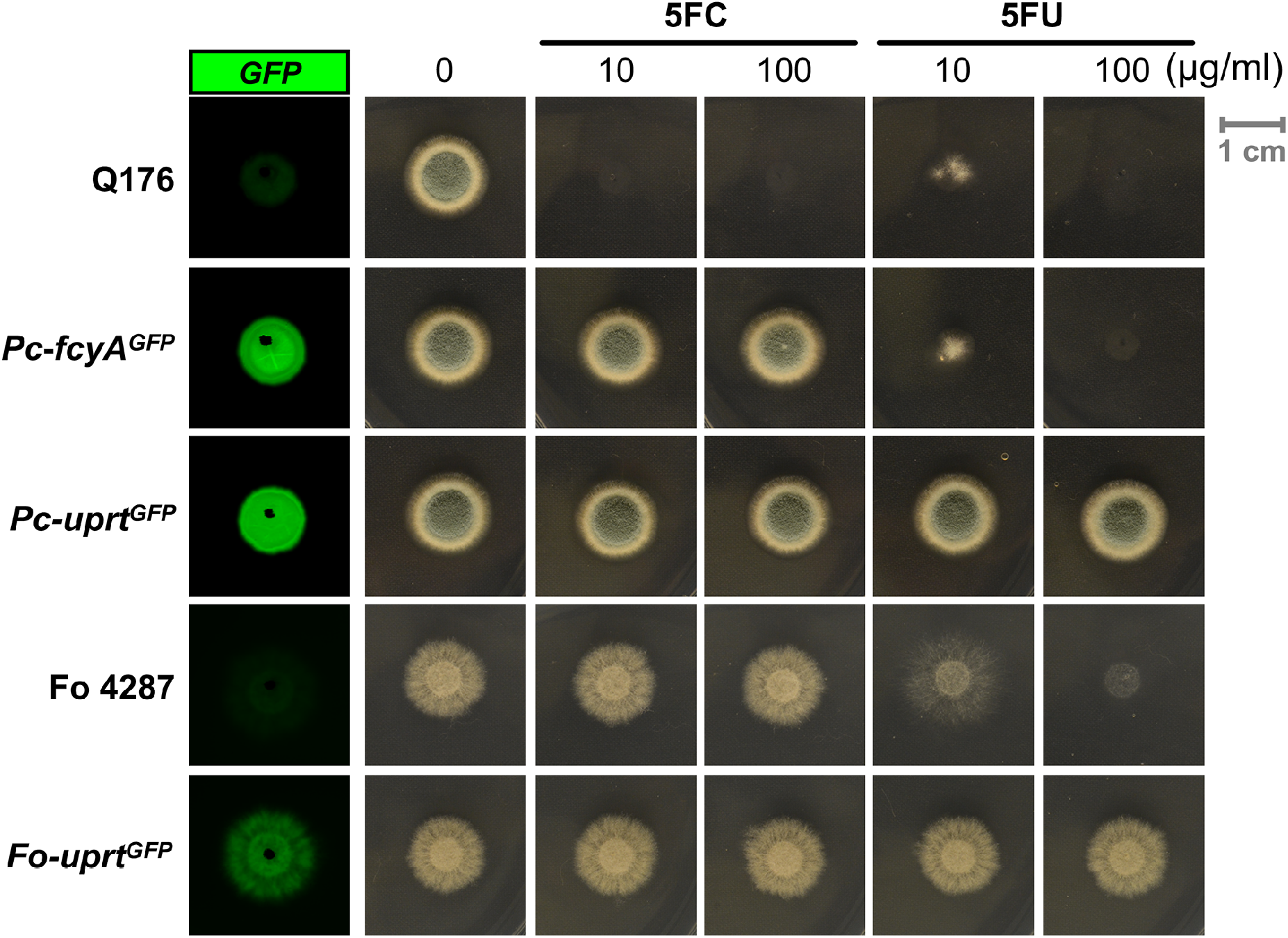
*P. chrysogenum* and *F. oxysporum* strains with replaced pyrimidine salvage pathway encoding loci. GFP expression was visualized in the corresponding knock-in mutants in the absence of drugs. 5FC/5FU resistance phenotypes of GFP reporter strains lacking CD *(Pc-fcyA^GFP^)* or UPRT *(Pc-uprt^GFP^, Fo-uprt^GFP^)* activities. For both experiments, strains were grown on solid AMM (*P. chrysogenum)* and PDA (*F. oxysporum)* followed by 72 h incubation at 25 °C.

Taken together, these data indicate high evolutionary conservation of the described pyrimidine salvage enzymes within the fungal clade and demonstrate the suitability of the encoding loci as markers for transformation selection in *P. chrysogenum* and *F. oxysporum*.

## DISCUSSION

In this study, we report the characterization and exploitation of multiple, endogenously-encoded selectable markers for targeted genetic engineering. The marker genes encode activities mediating 5FC uptake *(fcyB)* and metabolization *(fcyA* and *uprt)* of 5FC or 5FU into cell-toxic nucleotides (Fig. 1a) (Vermes et al. 2000). Genomic replacement of these genes by DOIs results in loss of the corresponding activities and can therefore be selected *via* 5FC or 5FU resistance. We validated the applicability of orthologous marker genes encoding pyrimidine salvage activities for their use as selectable markers in three fungal species (*A. fumigatus*, *P. chrysogenum* and *F. oxysporum*) by targeted insertion of various fluorescent or enzymatic reporter genes (Fig. 2, 3 and 5). Mimicking a biotechnological application, we further introduced the 17-kb penicillin biosynthetic gene cluster from *P. chrysogenum* to *A. fumigatus*, which naturally is not capable of producing this secondary metabolite. In addition to their single use, we demonstrate the possibility of consecutive use of these markers in a single strain by generating an *A. fumigatus* mutant expressing three fluorescent proteins (mKate2, sGFP and mTagBFP2) targeted to different subcellular compartments (Fig. 3a and Supplementary Fig. 4). Importantly, these endogenous genes can be used in addition to traditional selectable markers and the absence of all three genes (Δ*fcyB*Δ*fcyA*Δ*uprt*) affected neither growth nor virulence (Fig. 3b and Supplementary Fig. 5) in *A. fumigatus*, which represents a prerequisite for downstream use of engineered strains. Inspection of genomic sequences combined with 5FC/5FU susceptibility profiling of different fungal species playing important roles in biotechnology, medicine or agriculture revealed the presence of *uprt* orthologs in 10 out of 10 species, and *fcyB* as well as *fcyA* orthologs in 8 out of 10 species (Supplementary Table 1), indicating the broad applicability of the proposed selection method. Most likely, the number of self-encoded selectable markers for DOI integration allows expansion beyond the here characterized genes. For instance, genes coding for 5FU uptake, such as orthologs of *A. nidulans furD* or *S. cerevisiae FUR4* (Jund and Lacroute 1970; Hamari et al. 2009), represent potential candidates.

Employing pyrimidine salvage pathway-based endogenous markers, virtually any DOI can be site-specifically inserted into the target genome, demonstrating their huge potential for diverse applications in basic as well as applied research. The sequential use of marker genes enables the establishment of modular toolboxes, e.g. allowing site-directed insertion of multiple reporter genes for co-localization studies. Furthermore, the possibility to equip strains with multiple DNA building blocks makes this technology particularly attractive for synthetic biology. Notably, the fact that marker genes have to be inactivated to allow selection based on 5FC/5FU resistance enforces targeted integration of DOIs. As the described strategy avoids the use of foreign selection markers, the described genetic toolset further enables so called ‘self-cloning’. Strains engineered this way are not considered genetically modified organisms (GMO) in some countries (Steensels et al. 2014; Borner et al. 2019). For this, only endogenously encoded DOIs can be used; examples include the increase of gene dosage or engineering of sequences such as promoter swapping. A further advantage of self-encoded markers is the dispensability of antibiotic resistance genes, the use of which is generally discouraged in the manufacturing of medical- or food-related products (Yau and Stewart 2013; Mignon et al. 2015) as it may promote horizontal transfer of resistance genes.

In summary, we characterized a powerful genetic toolbox enabling multiple, targeted genomic insertions of DOIs. Evolutionary conservation of the pyrimidine salvage pathway suggests broad applicability of the described marker genes. Thus, we anticipate that this technology will significantly advance genetic and metabolic engineering of diverse organisms.

## MATERIALS AND METHODS

### Growth conditions and fungal transformation

Plate growth assays analyzing *A. fumigatus* and *P. chrysogenum* were carried out using solid *Aspergillus* minimal medium (AMM; ammonium tartrate was used as nitrogen source, glucose as carbon source)(Pontecorvo et al. 1953), and for *F. oxysporum* solid PDA was employed. Therefore, 10^4^ spores of each strain were point inoculated on agar plates. Low pH medium contained 100 mM citrate buffer (pH5), and neutral pH medium 100 mM MOPS buffer (pH7). If not stated in the text or images, plate growth assays were performed at pH5. For strains carrying *xylP* promoter (*PxylP*) driven reporter genes (*sGFP*, *mKate2^PER^*, *sGFP^MIT^*, *lacZ*), the medium was supplemented with 0.5% xylose to induce gene expression.

For fungal manipulations, 2 μg DNA of each construct was transformed into protoplasts of the respective recipient. For the regeneration of transformants, solid AMM (*A. fumigatus* and *P. chrysogenum)* or PDA (*F. oxysporum)* supplemented with 342 g/l or 200 g/l sucrose, respectively, were used. Selection procedures using conventional selectable marker genes (*hph*, *ble*) were carried out as described previously for *A. fumigatus* (Gsaller et al. 2018).

### Deletion of *A. fumigatus fcyA* and *uprt*

Strains and primers used in this study are listed in Supplementary Table 3 and 4. Coding sequences of *fcyA* and *uprt* were disrupted in wt (A1160P+) using hygromycin B and zeocin resistance cassettes, respectively. Therefore, deletion constructs comprising approximately 1 kb of 5’ and 3’ NTR linked to the central antibiotic resistance cassette were generated using fusion PCR as previously described (Fraczek et al. 2013). Correct integration of constructs was confirmed by Southern analyses (Supplementary Fig. 1).

### Generation of *A. fumigatus* knock-in strains

Knock-in constructs for *A. fumigatus* loci *fcyB, fcyA* and *uprt, P. chrysogenum* loci *Pc-fcyA* and *Pc-uprt*, as well as *F. oxysporum Fo-uprt* were generated similarly to the gene deletion fragments described above using fusion PCR. Here, instead of the antibiotic resistance cassettes, DOIs (for reporter templates, see also Supplementary Fig. 7) were connected to approximately 1 kb 5’ and 3’ NTR of the respective locus (Fig. 2a).

### LacZ-based colorimetric assay and fluorescence imaging

For the detection of LacZ activity (conversion of X-Gal into the blue compound 5,5’-dibromo-4,4’-dichlor-indigo) (Horwitz et al. 1964), a 5 ml layer of a 1 mM X-Gal/1% agar/1% N-lauroylsarcosin solution was poured over fungal colonies. GFP expression of fungal colonies was visualized using the laser scanner Typhoon FLA9500 (Ex 473 nm; Em ≥ 510 nm).

Images of *RFP^PER^GFP^MIT^BFP^CYT^* were taken using an HC PL APO CS2 63x/1.30 glycerol objective on an SP8 confocal microscope (Leica Microsystems, Wetzlar, Germany) equipped with a 80 Mhz pulsed white light laser (WLL) and a 405 nm CW diode laser (405 nm diode) according to Nyquist sampling. Gating of the red signal only was used in order to remove unspecific red autofluorescence. Images of mKate2^PER^ (Ex: 588 nm WLL, Em: 598-750 nm, Gating: 0.2. to 8 ns), sGFP^MIT^ (Ex: 489 nm WLL, Em: 499 – 578 nm) and mTagBFP^CYT^ (Ex: 405 nm Diode, Em: 415 – 479 nm) were processed using ImageJ (Fig. 3a). For the generation of 3D reconstructions (Supplementary Fig. 4), z-stacks, acquired under the same imaging conditions using a z-interval of 180 nm, were deconvolved using the CMLE algorithm of Huygens Professional version 18.10 (Scientific Volume Imaging, Amsterdam, The Netherlands) and further processed with Imaris 9.3.0 (Bitplane, Zurich, Switzerland).

### Expression analysis of penicillin biosynthetic genes and detection of penicillin G in culture supernatants

Northern analysis was carried out as described previously, using digoxigenin-labeled probes (Gsaller et al. 2012).

To detect the potential production of penicillin G, strains were grown in AMM for 48 h at 25 °C and 2 ml of culture supernatant were extracted with 1 volume of butyl acetate. The organic phase was collected in a new reaction tube and dried using a centrifugal vacuum concentrator (speed-vac). NanoLC-MS-based detection of penicillin G was carried out using an UltiMate 3000 nano-HPLC system coupled to a Q Exactive HF mass spectrometer (Thermo Scientific, Bremen, Germany). The samples were separated on a homemade fritless fused-silica microcapillary column (100 μm i.d. x 280 μm o.d. x 19 cm length) packed with 2,4 μm reversed-phase C18 material (Reprosil). Solvents for HPLC were 0.1% formic acid (solvent A) and 0.1% formic acid in 85% acetonitrile (solvent B). The gradient profile was as follows: 0-4 min, 4% B; 4-57 min, 43-5% B; 57-62 min, 35-100% B, and 62-67 min, 100% B. The flow rate was 300 nL/min.

Mass spectra were acquired in positive ion mode applying a precursor scan over the m / z range 50 to 500 in the FT analyzer. The ions at m / z = 335.1060 were selected from this precursor scan for MS/MS fragmentation in the linear ion trap.

### Murine infection model

Specific-pathogen-free female outbreed CD-1 mice (18–20 g, 6–8 weeks old, Charles River, Germany) were housed under standard conditions in individually ventilated cages and supplied with normal mouse chow and water *ad libitum*. All animals were cared for in accordance with the European animal welfare regulations, and experiments were approved by the responsible federal/state authority and ethics committee in accordance with the German animal welfare act (permit no. 03-027/16). Mice were immunosuppressed with cortisone acetate and intranasally infected with 2 x 10^5^ conidia in 20 μl PBS as described before (McCormick et al. 2012). Infected animals were monitored twice daily and humanely sacrificed if moribund (defined by a score including weight loss, piloerection, behavior and respiratory symptoms).

## ACKNOWLEDGMENTS

This project received funding from the Austrian Science Fund (FWF) P31093-B32 and J3651-B22 to FG. MS L-B. was supported by the FWF through the ‘Lise Meitner Program’ (M1962-B21).

## AUTHOR CONTRIBUTIONS

LB, AD, MS L-B, MO, MS, BS, OS, IB, BA and FG performed experiments; LB, AD, MS L-B, IDJ, MS, MO, HL, BS, HH and FG analyzed data; LB, IDJ, HL, HH and FG designed experiments; LB, HH and FG wrote the manuscript; FG conceptualized the study; all authors edited the manuscript.

