## Supplementary Information for "Multiplex genetic engineering exploiting pyrimidine salvage pathway-based self-encoded selectable markers"

†, Present address:
Department of Genetics, University of Córdoba, 14071 Córdoba, Spain

**SUPPLEMENTARY TABLES**

Supplementary Table 1 **Homologous proteins to *A. fumigatus* (A1163) FcyB, FcyA and Uprt in fungal species with relevance in biotechnology, agriculture or medicine identified by BLASTP analysis (https://blast.ncbi.nlm.nih.gov/Blast.cgi).** The best hits, showing > 40% identity, are illustrated.

| FcyB | Species | Total score | Query coverage | E-value | Identity | Protein Accession |
|---|---|---|---|---|---|---|
| | *Aspergillus fumigatus A1163* | 1060 | 100% | 0 | 100% | EDP54513.1 |
| | *Aspergillus oryzae RIB40* | 855 | 100% | 0 | 80% | XP_001826247.1 |
| | *Aspergillus niger ATCC 1015* | 818 | 100% | 0 | 77% | EHA22089.1 |
| | *Penicillium chrysogenum* | 780 | 99% | 0 | 75% | KZN90676.1 |
| | *Saccharomyces cerevisiae P283* | 415 | 98% | 2.00E-139 | 42% | EWH18811.1 |
| | *Candida albicans SC5314* | 411 | 91% | 2.00E-138 | 46% | XP_714531.2 |
| | *Saccharomyces cerevisiae P283* | 411 | 98% | 3.00E-138 | 43% | EWH18815.1 |
| | *Komagataella phaffii GS115* | 395 | 99% | 3.00E-132 | 42% | XP_002493949.1 |
| | *Cryptococcus neoformans var. grubii H99* | 320 | 98% | 5.00E-103 | 40% | XP_012052683.1 |
| **FcyA** | **Species** | **Total score** | **Query coverage** | **E-value** | **Identity** | **Protein Accession** |
| | *Aspergillus fumigatus A1163* | 303 | 100% | 3.00E-107 | 100% | EDP55842.1 |
| | *Aspergillus niger ATCC 1015* | 280 | 100% | 4.00E-98 | 91% | EHA26383.1 |
| | *Penicillium chrysogenum* | 278 | 100% | 2.00E-97 | 91% | KZN93743.1 |
| | *Aspergillus oryzae RIB40* | 275 | 95% | 4.00E-96 | 93% | XP_001819938.3 |
| | *Komagataella phaffii GS115*[#] | 196 | 95% | 1.00E-64 | 63% | XP_002490927.1 |
| | *Candida albicans SC5314* | 189 | 95% | 3.00E-62 | 61% | KHC73214.1 |
| | *Saccharomyces cerevisiae P283* | 181 | 95% | 5.00E-59 | 61% | EWH15533.1 |
| | *Cryptococcus neoformans var. grubii H99* | 140 | 93% | 2.00E-42 | 49% | XP_012046842.1 |
| **Uprt** | **Species** | **Total score** | **Query coverage** | **E value** | **Identity** | **Protein Accession** |
| | *Aspergillus fumigatus A1163* | 496 | 100% | 1.00E-180 | 100% | EDP51298.1 |
| | *Aspergillus niger ATCC 1015* | 464 | 99% | 4.00E-168 | 94% | EHA22482.1 |
| | *Penicillium chrysogenum* | 450 | 99% | 3.00E-162 | 89% | KZN87537.1 |
| | *Aspergillus oryzae RIB40* | 445 | 100% | 3.00E-160 | 90% | XP_023088768.1 |
| | *Trichoderma reesei QM6a* | 397 | 98% | 4.00E-141 | 78% | XP_006967593.1 |
| | *Fusarium oxysporum f. sp. lycopersici 4287* | 389 | 95% | 6.00E-139 | 78% | XP_018234120.1 |
| | *Komagataella phaffii GS115*[#] | 301 | 87% | 7.00E-104 | 67% | XP_002489914.1 |
| | *Cryptococcus neoformans var. grubii H99* | 301 | 85% | 1.00E-103 | 69% | XP_012050086.1 |
| | *Candida albicans SC5314* | 298 | 90% | 2.00E-102 | 66% | XP_712023.1 |
| | *Saccharomyces cerevisiae P283* | 294 | 87% | 4.00E-101 | 66% | EWH18153.1 |

[#] *old species name Pichia pastoris* (Heistinger et al. 2018).

Supplementary Table 2 **Susceptibility testing of fungal species.** MICs of 10 fungal species were determined following EUCAST guidelines (Subcommittee on Antifungal Susceptibility Testing of the 2008). All strains were uniformly incubated in RPMI at 30 °C for 48 h followed by visual assessment of MICs. A combination of BLAST based *in silico* analysis and susceptibility testing further suggested the presence or absence of *A. fumigatus* FcyB, FcyA or Uprt orthologs.

| | **MIC (µg/ml)** | | | | **Activities** | | |
|---|---|---|---|---|---|---|---|
| | **5FC** | | **5FU** | | | | |
| **Species** | **pH5** | **pH7** | **pH5** | **pH7** | **FcyB** | **FcyA** | **Uprt** |
| *Aspergillus fumigatus* | 0.39 | 400 | 50 | 100 | ✔ | ✔ | ✔ |
| *Aspergillus niger* | 0.39 | 6.25 | 50 | 100 | ✔ | ✔ | ✔ |
| *Aspergillus oryzae* | 0.39 | 400 | 200 | 400 | ✔ | ✔ | ✔ |
| *Candida albicans* | 0.39 | 0.39 | 50 | 100 | ✔ | ✔ | ✔ |
| *Cryptococcus neoformans* | 0.39 | 25 | 6.25 | 6.25 | ✔ | ✔ | ✔ |
| ***Fusarium oxysporum*** | **>400** | **>400** | 400 | 400 | | | ✔ |
| *Komagataella phaffii* | 0.39 | 0.39 | 0.39 | 0.39 | ✔ | ✔ | ✔ |
| *Penicillium chrysogenum* | 0.39 | 3.12 | 3.12 | 3.12 | ✔ | ✔ | ✔ |
| *Saccharomyces cerevisiae* | 0.39 | 0.39 | 0.8 | 50 | ✔ | ✔ | ✔ |
| ***Trichoderma reesei*** | **>400** | **>400** | 100 | 100 | | | ✔ |

**Bold**, strains lacking susceptibility to 5FC under both tested conditions, suggesting the absence of CD activity.
✔, indicates the presence of the corresponding activity.

Supplementary Table 3 **Strains used in this study.**

| Strain | Genotype | Reference |
|---|---|---|
| ***A. fumigatus*** | | |
| **Characterization of 5FC activity determinants** | | |
| A1160P+ (wt) | | (Fraczek et al. 2013) |
| Δ*fcyB* | Δ*fcyB::hph* | (Gsaller et al. 2018) |
| Δ*fcyA* | Δ*fcyA::hph* | This study |
| Δ*uprt* | Δ*uprt::ble* | This study |
| **Proof-of-principle work in *A. fumigatus*** | | |
| $fcyB^{GFP}$ | Δ*fcyB::PxylPsGFP* | This study |
| $fcyB^{lacZ}$ | Δ*fcyB::PxylPlacZ* | This study |
| $fcyA^{GFP}$ | Δ*fcyA::PxylPsGFP* | This study |
| $fcyA^{lacZ}$ | Δ*fcyA::PxylPlacZ* | This study |
| $uprt^{GFP}$ | Δ*uprt::PxylPsGFP* | This study |
| $uprt^{lacZ}$ | Δ*uprt::PxylPlacZ* | This study |
| **Multicolor laser scanning microscopy** | | |
| $RFP^{PER}$ | Δ$fcyB$::$mKate2^{PER}$ | This study |
| $RFP^{PER}GFP^{MIT}$ | Δ$fcyB$::$mKate2^{PER}$Δ$fcyA$::$sGFP^{MIT}$ | This study |
| $RFP^{PER}GFP^{MIT}BFP^{CYT}$ | Δ$fcyB$::$mKate2^{PER}$Δ$fcyA$::$sGFP^{MIT}$Δ$uprt$::$mTagBFP2^{CYT}$ | This study |
| **Introduction of the penicillin G biosynthetic cluster** | | |
| $fcyB^{PENG}$ | Δ*fcyB::PcCluster* | This study |
| **Implementation of the 5FC/5FU selection method in *P. chrysogenum* and *F. oxysporum*** | | |
| ***P. chrysogenum*** | | |
| Q176 (wild-type) | | (Backus and Stauffer 1955) |
| $Pc\text{-}fcyA^{GFP}$ | Δ*Pc-fcyA::PxylPsGFP* | This study |
| $Pc\text{-}uprt^{GFP}$ | Δ*Pc-uprt::PxylPsGFP* | This study |
| ***F. oxysporum*** | | |
| Fol 4287 (wild-type) | | (Ma et al. 2010) |
| $Fo\text{-}uprt^{GFP}$ | Δ*Fo-uprt::PgpdAGFP* | This study |
| **Strains used for the determination of MICs** | | |
| *A. niger CBS 120-49* | | (Bos et al. 1988) |
| *A. oryzae NS4* | | (Jin et al. 2004) |
| *C. albicans SC5314* | | (Maestrone and Semar 1968) |
| *C. neoformans H99* | | (Perfect et al. 1980) |
| *K. phaffii KM71* | | (Salamin et al. 2010) |
| *S. cerevisiae BY4742* | | (Brachmann et al. 1998) |
| *T. reesei QM9414* | | (Ghose and Sahai 1979) |

Supplementary Table 4 **Oligonucleotides used in this study.**

| primer name | forward primer (5´→3´) | reverse primer (5´→3´) | PCR product |
|---|---|---|---|
| ***(A) fcyA and uprt deletion and knock-in cassettes.*** | | | |
| fcyA-1/-2 | TTGAAACTCCGAGGAAGTCG | TAGTTCTGTTACCGAGCCGGTATGTGGATCCAGAGCGTCA | 5' fcyA |
| fcyA-3/-4 | GCTCTGAACGATATGCTCCCTTCGACAAAATGCCATTGAA | TACCTCCCCGAATACCATGA | 3' fcyA & 3' probe for Southern analysis |
| fcyA-N1/-N2 | CGAGTCGCCTTAAAATGAGC | GTGGATCGGTATGCAGGATT | fcyA::hph deletion; knock-in constructs |
| uprt-1/-2 | GGAAGGACAGGTACGCCATA | TAGTTCTGTTACCGAGCCGGCGGAGCACTCTGAAAATTGG | 5' uprt |
| uprt-3/-4 | GCTCTGAACGATATGCTCCCTCCCATCGTGTAGCGACATA | TACTACCTTCGCCCTCTGGA | 3' uprt & 3' probe for Southern analysis |
| uprt-N1/-N2 | TTTGAGCGATTAAGGTGCAA | GCCCCACTACTTGTTTCCAG | uprt::ble deletion; knock-in constructs |
| fcyB-3/-4 | GCTCTGAACGATATGCTCCCTGCGGTTTTTGGGTTTTATC | CACACTGGGTCTGAAGACGA | 3' probe for Southern analysis |
| ***(B) Amplification of reporter cassettes.*** | | | |
| P1/P2 | CCGGCTCGGTAACAGAACTACTGATGCGAGCAACAGTATGC | GGGAGCATATCGTTCAGAGCTGAGGGTTGAGTACGAGATTGG | reporter cassettes: sGFP*, lacZ, mKate2$^{PER}$, sGFP$^{MIT}$ |
| hph-FW/-RV | CCGGCTCGGTAACAGAACTAACGGCGTAACCAAAAGTCAC | GGGAGCATATCGTTCAGAGCTCTTGACGACCGTTGATCTG | hygR & zeoR cassette; mTagBFP2$^{CYT}$ reporter casse |
| FoGFP-FW/-RV | CGAGACCTAATACAGCCCCTA | CCTGTGCATTCTGGGTAAACG | GFP reporter cassette for *F. oxysporum* |
| * same cassette was used for the *P. chrysogenum* reporter | | | |
| ***(C) Generation of the PcCluster containing knock-in plasmid*** | | | |
| 5' fcyB-FW/-RV | TGTGGCGGCCGCGTTTAAACCGCTATCCCAGCAATAGAGC | TTACGCCAAGCTTGCATGCCACTGAGTCAATCCCCACCAC | 5' fcyB |
| 3' fcyB-FW/-RV | AGTGAATTCGAGCTCGGTACTGCGGTTTTTGGGTTTTATC | AGCGGTTTAAACGCGGCCGCCACACTGGGTCTGAAGACGA | 3' fcyB |
| BB-pfcyB-FW/-RV | TGTGAAATTGTTATCCGCTCACAA | AAACAGCTATGACCATGATTACGC | backbone pfcyB |
| PcFrag1-FW/-RV | AATCATGGTCATAGCTGTTTAAAGGGGAGAGAGCGAAAAG | GCATGGGGACAATCTCACTT | fragment 1 PcCluster |
| PcFrag2-FW/-RV | AAGTGAGATTGTCCCCATGCAG | GAGCGGATAACAATTTCACACGCGTGATATCCTGTCTTCA | fragment 2 PcCluster |
| ***(D) Generation of the knock-in constructs for P. chrysogenum and F. oxysporum*** | | | |
| Pc-fcyA-1/-2 | TGACCTTGATGGCATCTGAA | TAGTTCTGTTACCGAGCCGGTCAGTGCGGGCTACAGAGTA | 5' Pc-fcyA & 5' probe for Southern analysis |
| Pc-fcyA-3/-4 | GCTCTGAACGATATGCTCCCGGCCTGCACATATCATAGCC | AGCCGTAAAATTCGCATCAC | 3' Pc-fcyA |
| Pc-fcyA-N1/-N2 | GTCGAGGTGCTCAATGTGAA | TTGTTTTGACTTCCCCTTCG | Pc-fcyA knock-in construct |
| Pc-uprt-1/-2 | GGACAGTTTGGACAATGCAG | TAGTTCTGTTACCGAGCCGGTTTGAAGGGCAAGAGTCCAG | 5' Pc-uprt & 5' probe for Southern analysis |
| Pc-uprt-3/-4 | GCTCTGAACGATATGCTCCCACCACGTTGAAAGGAGCATC | AGACCGTGGAAGTTGGTCAG | 3' Pc-uprt |
| Pc-uprt-N1/-N2 | TTTTGCAAGGGTCGAGAAAG | CAGTTCTTGCCCTGGATCTC | Pc-uprt knock-in construct |
| Fo-uprt-1/-2 | CATACGTCACCACCTTGC | GTTGTAGGGGCTGTATTAGGTCTCGGCTGTTGTTAGTGTTCGAGG | 5' Fo-uprt & 5' probe for Southern analysis |
| Fo-uprt-3/-4 | GAGTCGTTTACCCAGAATGCACAGGGAAGGAATCAGCGCAAAG | CACGTATAGAATCACGGAGG | 3' Fo-uprt |
| Fo-uprt-N1/-N2 | GACGCCATAGTGTGCTC | GCTTGATGCATGCACTAG | Fo-uprt knock-in construct |

### SUPPLEMENTARY FIGURES

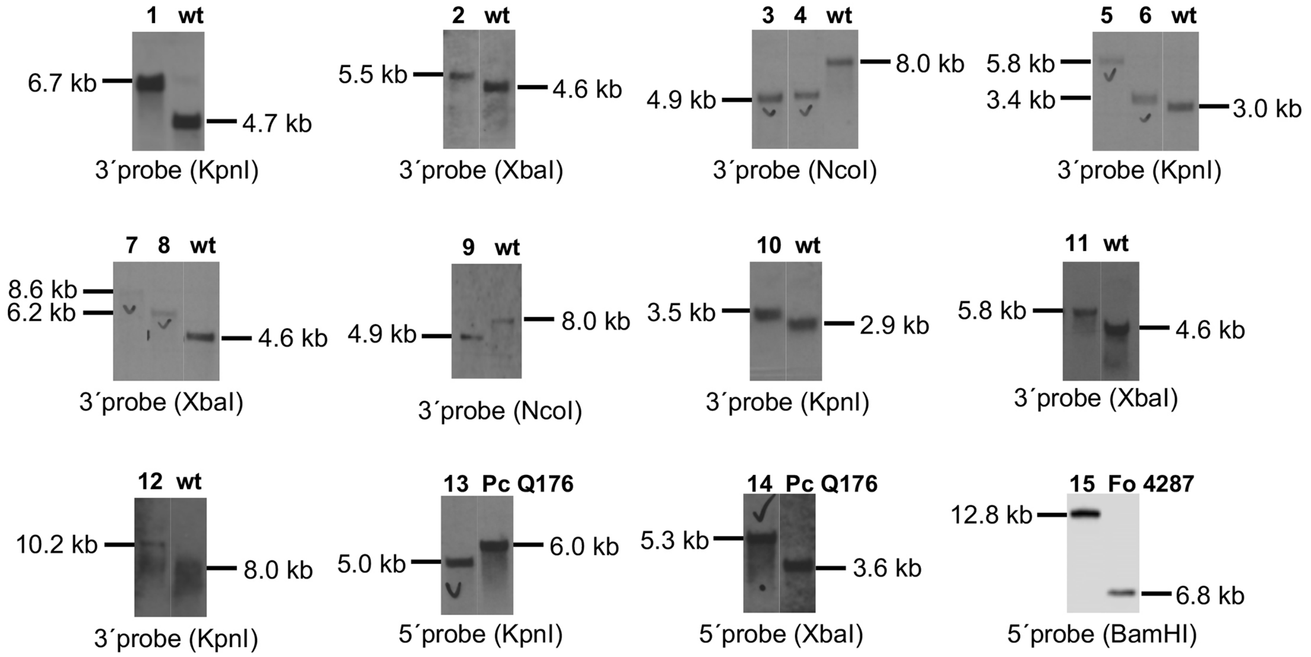

1, Δ*fcyA::hph;* 2, Δ*uprt::ble;* 3, Δ*fcyB::PxylPlacZ* ; 4, Δ*fcyB::PxylPsGFP* ; 5, Δ*fcyA::PxylPlacZ* ; 6, Δ*fcyA::PxylPsGFP* 7, Δ*uprt::PxylPlacZ* ; 8, Δ*uprt::PxylPsGFP;* 9, Δ*fcyB*::*mKate2*$^{PER;}$ 10, Δ*fcyB*::*mKate2*$^{PER}$Δ*fcyA*::*sGFP*$^{MIT}$ ; 11, Δ*fcyB*::*mKate2*$^{PER}$Δ*fcyA*::*sGFP*$^{MIT}$Δ*uprt*::*mTagBFP2*$^{CYT;}$ 12, Δ*fcyB*::*PcCluster;* 13, Δ*Pc-fcyA*::*PxylPsGFP;* 14, Δ*Pc-uprt*::*PxylPsGFP;* 15, Δ*Fo-uprt*::*PgpdAGFP*

Supplementary Fig. 1 **Southern blot analysis of strains generated in this work.** In each blot a representative transformant is compared to the respective recipient strain; wt, *A. fumigatus* A1160P+; Pc, *Penicillium chrysogenum*, Fo, *Fusarium oxysporum*.

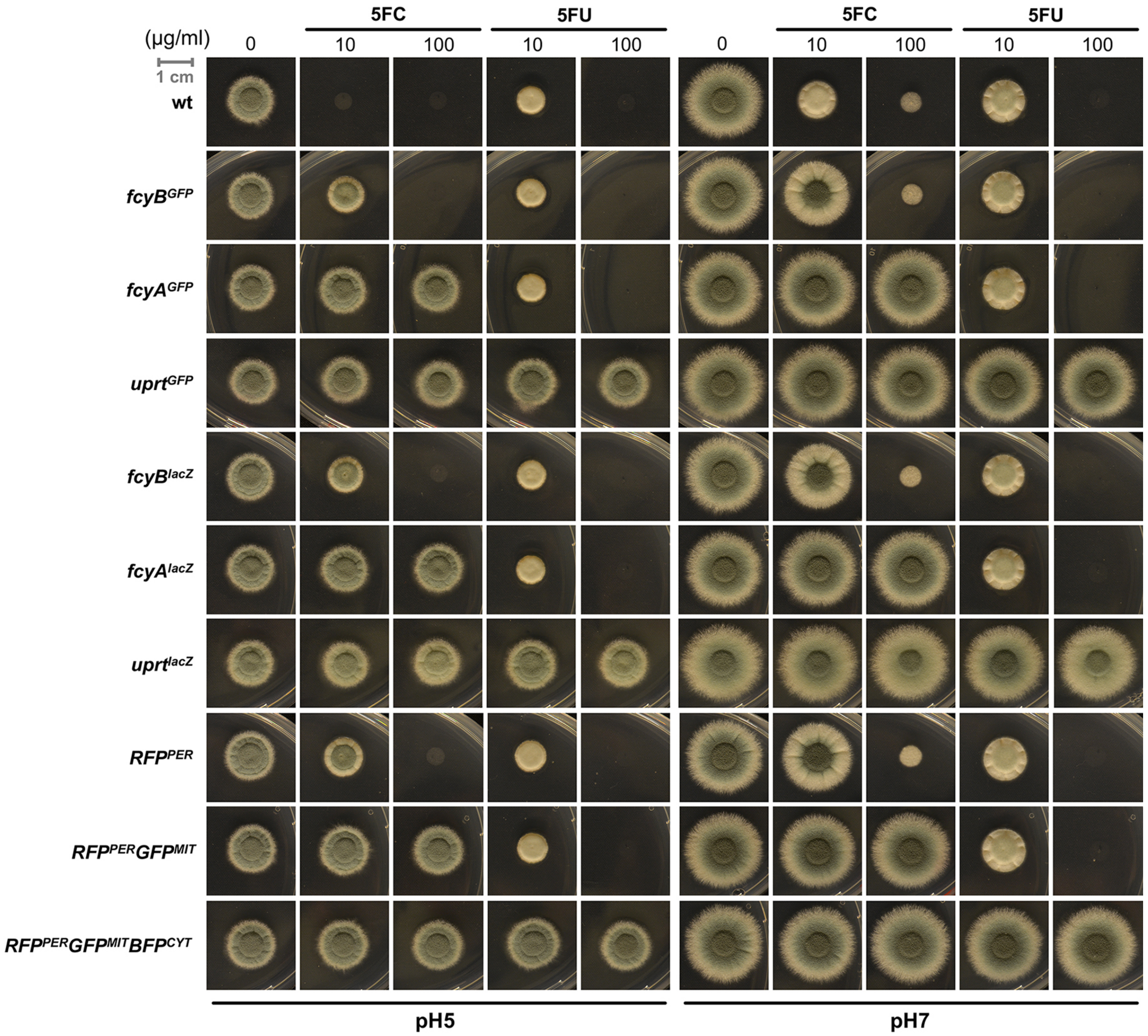

Supplementary Fig. 2 **Plate growth based 5FC/5FU susceptibility testing of *A. fumigatus* GFP and LacZ knock-in strains as well as *$RFP^{PER}GFP^{MIT}BFP^{CYT}$* and its progenitor strains.** Strain were point inoculated on solid AMM at both pH5 and pH7 and incubated for 48 h at 37°C. Resistance phenotypes of all mutants analyzed were in accordance with the absence of individual salvage activities.

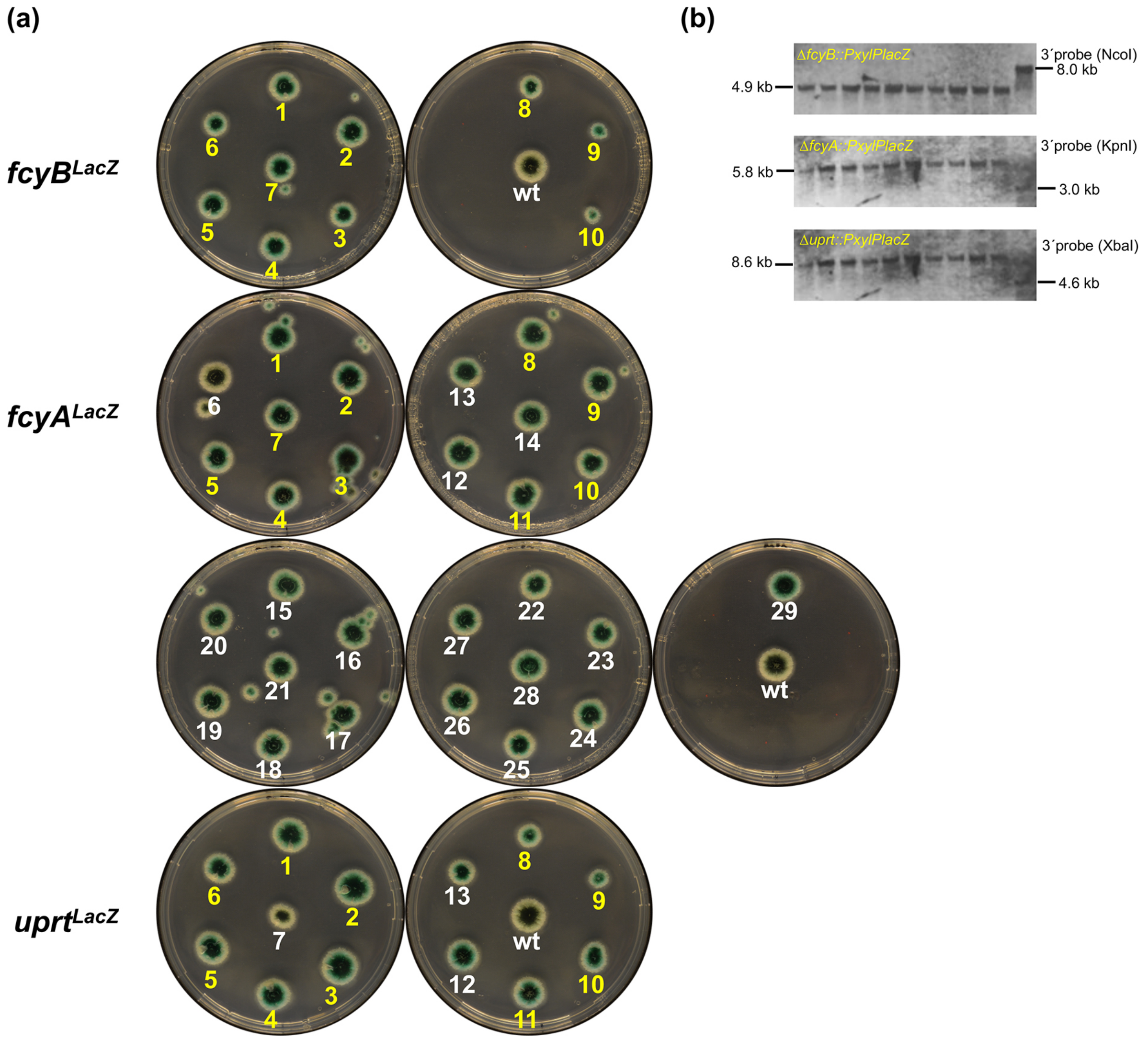

Supplementary Fig. 3 **β-galactosidase staining to screen for LacZ-positive transformants.** After determining LacZ activities of each transformant (a), 10 transformants per locus showing LacZ-positive phenotypes (yellow numbers) were subject to Southern blot analysis (b). Strains were grown for 48 h at 37 °C on solid AMM before pouring an additional 5ml-layer of X-Gal containing agar on the top of colonies.

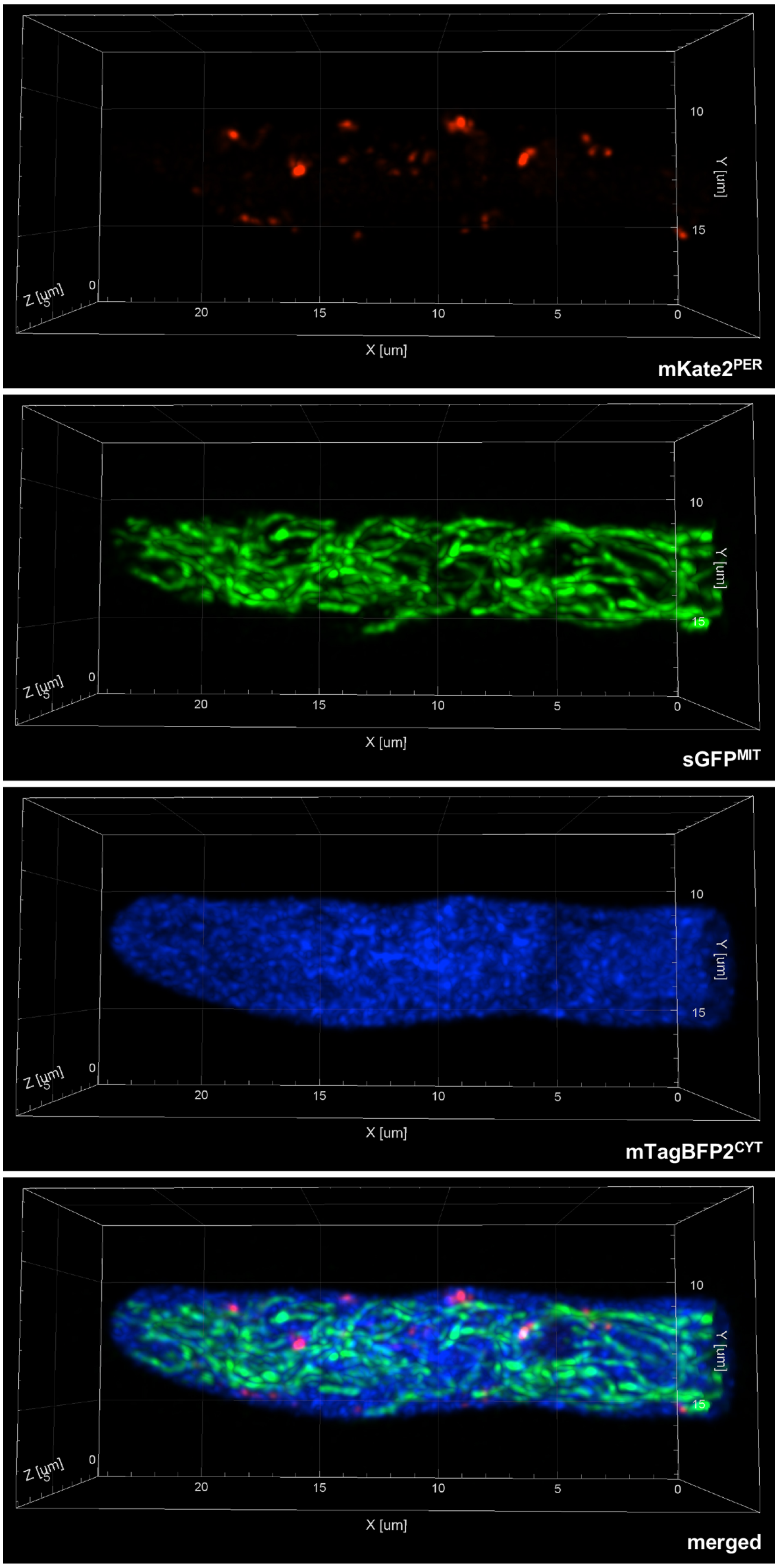

Supplementary Fig. 4 **Multicolor imaging of *$RFP^{PER}GFP^{MIT}BFP^{CYT}$*.** 3D reconstruction of single and merged channels for *$mKate2^{PER}$* (peroxisomal), *$sGFP^{MIT}$* (mitochondrial) and *$mTagBFP2^{CYT}$* (cytosolic).

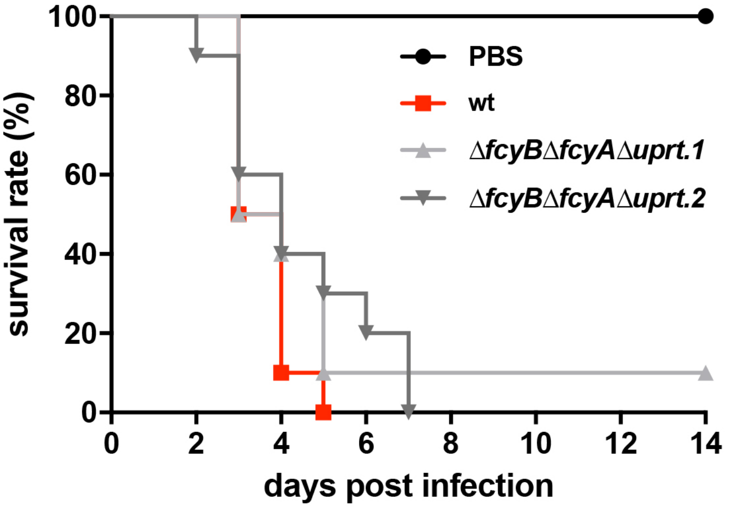

Supplementary Fig. 5 **Simultaneous lack of the three pyrimidine salvage pathway genes *fcyB*, *fcyA* and *uprt* does not affect *A. fumigatus* virulence in a pulmonary murine model of aspergillosis.** Survival of female outbreed CD-1 mice immunosuppressed with cortisone acetate and intranasally infected with 2 x $10^5$ conidia in 20 µl PBS is shown as Kaplan-Meier curves. *ΔfcyBΔfcyAΔuprt.1* and *ΔfcyBΔfcyAΔuprt.2* represent two independent knock-in transformants lacking FcyB, FcyA and Uprt. Analysis by log-rank test showed no significant differences ($p > 0.05$ in comparison to wt and comparison between mutants; mock infected n=5, infected animals n=10/group).

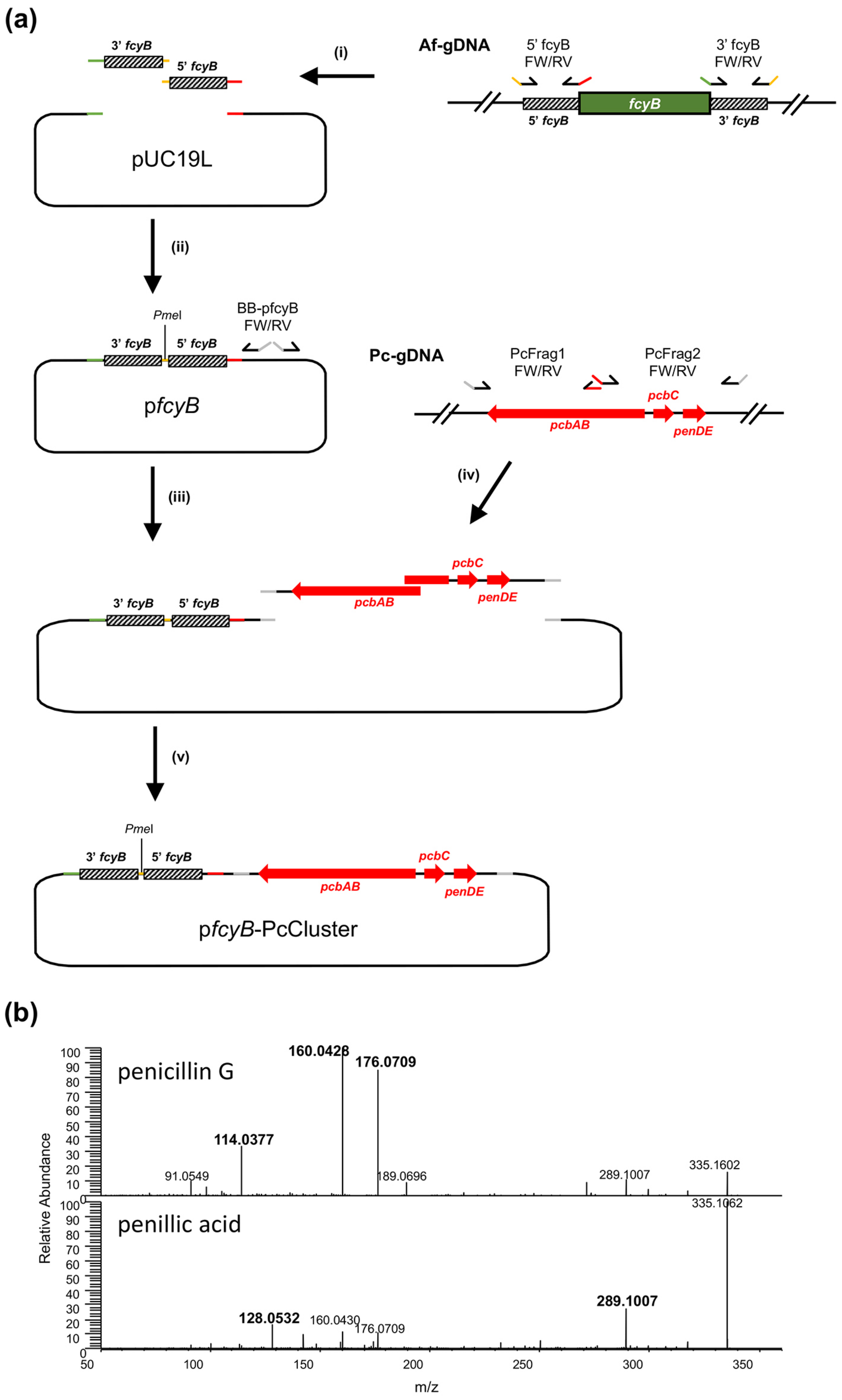

(a)
3' fcyB
5' fcyB
(i)
Af-gDNA
5' fcyB FW/RV
3' fcyB FW/RV
fcyB
pUC19L
(ii)
PmeI
BB-pfcyB FW/RV
pfcyB
Pc-gDNA
PcFrag1 FW/RV
PcFrag2 FW/RV
pcbC
pcbAB
penDE
(iii)
(iv)
(v)
pfcyB-PcCluster
(b)
penicillin G
160.0428
176.0709
114.0377
91.0549
189.0696
289.1007
335.1602
penillic acid
335.1062
289.1007
128.0532
160.0430
176.0709
Relative Abundance
m/z

Supplementary Fig. 6 **Generation of pfcyB-PcCluster and fragmentation patterns of penicillin G and penillic acid.** (a) After amplification of fcyB 5'- (5' fcyB-FW/-RV) and 3'-NTRs (3' fcyB-FW/-RV) from *A. fumigatus* genomic DNA (Af-gDNA) (i), the purified fragments were assembled (NEBuilder®) into pUC19L (ii). The primers 5' fcyB-FW and 3' fcyB-RV contained an add-on sequence including the *Pme*I restriction site. The yielding plasmid pfcyB was linearized by PCR amplification (iii) using primers BB-pfcyB-FW/RV. Two overlapping fragments comprising the penicillin G biosynthetic cluster were amplified from *P. chrysogenum* genomic DNA (Pc-gDNA) employing primer pairs PcFrag1-FW/RV and PcFrag2-FW/RV (iv). PcFrag1, PcFrag2 and linear pfcyB were assembled (v) giving rise to pfcyB-PcCluster. (b) Structure-specific fragmentation patterns of penicillin G (m/z 335.1060 -> 160.04, 176.07 and 114.04; upper panel) and penillic acid (m/z 335.1060 -> 289.10 and 128.05; lower panel) are illustrated (Aldeek et al. 2016).

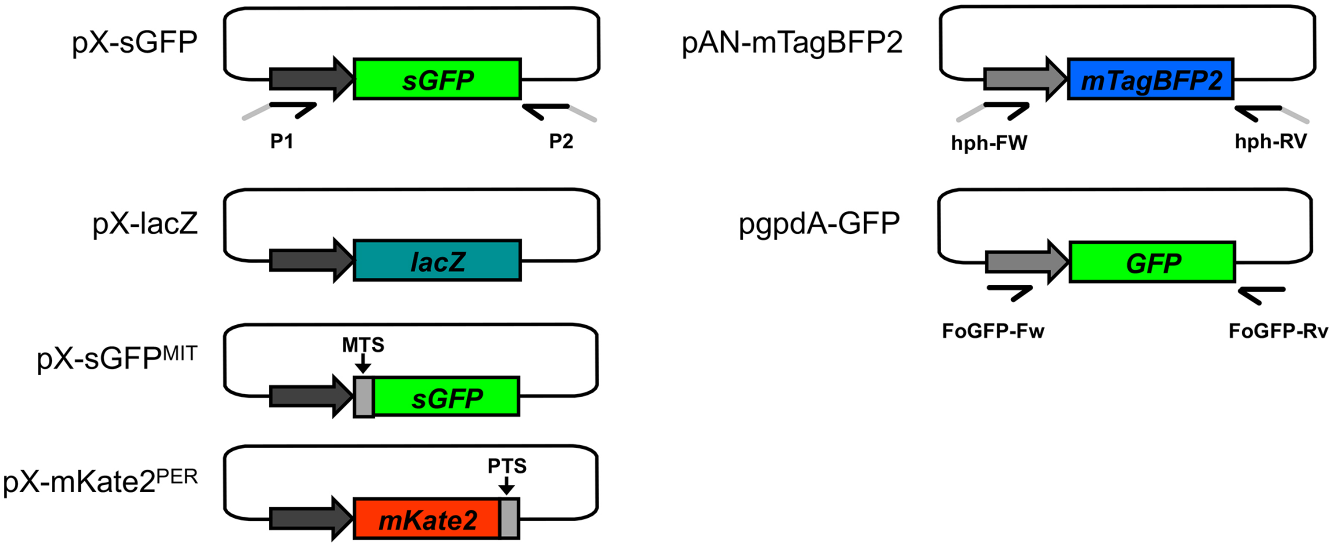

Supplementary Fig. 7 **Plasmid templates used for the generation of the different DOIs transformed in this work.** For the amplification of the reporter cassettes comprising *sGFP*, *lacZ*, *mKate2*$^{PER}$, *sGFP*$^{MIT}$ from the 'pX' plasmids pX-sGFP, pX-mKate2$^{PER}$, pX-sGFP$^{MIT}$, pX-lacZ the primer pair P1/P2 was used. An mTagBFP2 containing cassette was amplified from pAN-mTagBFP2 using primers hph-FW/hph-RV. For *F. oxysporum*, the GFP reporter cassette was amplified from pgpdA-GFP using primers FoGFP-Fw/Rv. In 'pX' plasmids, the reporter genes are under control of the xylose-inducible promoter *PxylP*; in the other two plasmids, the reporter genes are driven by the constitutive *gpdA* promoter derived from *A. nidulans* (Punt et al. 1988). Other abbreviations: MTS, mitochondrial targeting sequence; PTS, peroxisomal targeting sequence.
